# Biomechanics of yawning: insights into cranio-cervical fluid dynamics and kinematic consistency

**DOI:** 10.64898/2025.12.17.695005

**Authors:** Adam D. Martinac, Stean Waters, Robert A. Lloyd, Lynne E. Bilston

## Abstract

Yawning is a stereotyped orofacial–respiratory behaviour whose physiological role remains uncertain. Because cerebrospinal fluid (CSF) movement contributes to solute transport and waste removal and is strongly influenced by respiratory pressure dynamics, the study evaluated whether yawning alters neurofluid flow relative to normal and deep breathing, and whether yawning kinematics are reproducible within individuals. In a single MRI session in healthy adults, real-time phase-contrast imaging at the upper cervical level (C3) was combined with mid-sagittal real-time cine imaging to quantify CSF and internal jugular venous flows during normal breathing, forceful oral inspirations (deep breaths), yawns, and stifled yawns, and to derive tongue-motion trajectories. Both deep breaths and yawns increased CSF and venous flow compared with normal breathing; despite similar flow magnitudes, yawns more frequently produced co-directional caudal CSF and jugular outflow during inspiration, whereas deep breaths typically showed counter-directional CSF–venous flow. Yawning kinematics were highly reproducible within individuals across repeated events, indicating a stable motor sequence consistent with brainstem pattern-generator control. These observations show that yawning is not simply an intensified breath but a distinct cardiorespiratory manoeuvre that reorganizes neurofluid flow. The inspiratory alignment of CSF with venous outflow during yawns suggests a transient caudal advection that could influence solute transport and heat exchange within the cranial–cervical system, motivating targeted mechanistic studies with simultaneous airway pressure, thoraco-abdominal motion, and cervical venous pressure measurements.

## Introduction

Yawning, a respiratory manoeuvre characterised by mouth opening and deep inspiration, is a behaviour observed in a wide range of mammals, amphibians, reptiles, and other vertebrates (Moyaho et al., 2017; Palagi et al., 2019; Gallup and Wozny, 2022). Although yawning is a common behaviour, it remains poorly understood and experimental data on its mechanics are limited (Corey et al., 2012; Gupta & Mittal, 2013). Most yawns appear to be comprised of an initial deep inspiration, followed by a pause and then rapid expiration. A range of hypotheses have been proposed: physiological hypotheses suggest that yawning may play a role in regulating blood oxygen/CO₂ levels and airway patency (hypoxia/ventilation; Provine et al., 1987; Doelman & Rijken, 2022), brain thermoregulation (Gallup & Gallup, 2007; Gallup & Eldakar, 2013; Eldakar et al., 2015), arousal/attentional state (Guggisberg et al., 2007; Guggisberg et al., 2010; Thompson., 2014; Krestel et al., 2018), and cerebral metabolic waste clearance (Dolkart, 2017). In contrast, psychological hypotheses focus on the social and communicative aspects of yawning; see Massen & Gallup (2017).

CSF movement is critical for solute transport and metabolic waste removal, yet the influence of yawning on CSF and blood dynamics is largely unexplored. In general, CSF is thought to be produced by the choroid plexus in the brain’s ventricles, flowing through various pathways before exiting via the ventricular system and entering the subarachnoid space. The movement of the jaw and the act of inhaling can impact circulation within the skull. Specifically, any behaviour that compresses the jugular vein in the neck can immediately raise CSF pressure (Walusinski et al., 2013). Research by Lloyd et al. (2020) has shown that CSF flow is influenced by the pressures in the thoracic and lumbar spinal regions, which fluctuate during respiration, along with cranial and spinal blood flows. An increase in caudal CSF has been observed in the cervical spine and ventricles following the inspiratory phase of yawning, accompanied by an increase in caudal blood flow through the internal jugular vein (Schroth and Klose., 1992). This suggests that the physiological impacts of yawning might be reflected in the flow profiles of CSF and blood.

The highly stereotyped nature of yawns has led to the proposal that they are coordinated by a brainstem central pattern generator (CPG), like those controlling breathing and locomotion (Erkoyun et al., 2017). Because a brainstem CPG orchestrates a stereotyped, modulable sequence of inspiratory drive and orofacial–pharyngeal muscle activation, it could plausibly contribute to intrathoracic pressure transients and cervical venous pressure gradients that influence neurofluid (CSF and blood) flow.

Building on evidence that deep breathing strongly modulates blood and CSF flows in the cranium and spine (Yamada et al., 2013; Dreha-Kulaczewski et al., 2015, 2018; Lloyd et al., 2020; Kollmeier et al., 2022), our study investigates how yawning influences CSF and blood flow through the cervical spine. In parallel, this study aimed to examine the intra-individual reproducibility of yawning kinematics. Real-time phase-contrast magnetic resonance imaging (PC-MRI) and real-time sagittal scans were used as non-invasive methods to quantify both fluid velocities and anatomical movement during the respiratory manoeuvres. We hypothesised that the effect of yawning on CSF and blood flow would resemble the effect of a gaping deep breath. An additional hypothesis was that tongue motion patterns during repeated yawns would exhibit highly consistent, stereotyped patterns within individuals. Evidence of highly consistent stereotyped yawn-related tongue and jaw motion would add weight to existing evidence for the existence of a yawning CPG.

## Methods

### Ethics approval

All participants in this study provided informed, written consent. UNSW human research ethics committee (HREC ethics HC200474) approved the experimental protocols, and all experiments were conducted according to the approved protocol, and they were conducted according to the Declaration of Helsinki except for registration in a public database.

### Subjects

Twenty-two healthy participants with no contraindications for MRI were enrolled in the study, 11 males (age 31±11 years; weight 86±8 kg; height 178±6 cm; mean ± S.D.) and 11 females (age 37±17 years; weight 65±14 kg; height 174±9 cm; mean ± S.D.).

### Respiratory manoeuvres and breathing protocol

Participants performed four respiratory manoeuvres during MRI scanning: normal breathing, yawning, stifled yawning (voluntary suppression of a yawn), and a forceful submaximal oral inspiratory effort (hereafter referred to as a “deep breath”) designed to reproduce the jaw opening and airflow characteristics of a yawn. Prior to scanning, all participants were trained and given the opportunity to practise each manoeuvre. Yawns and stifled yawns could not be standardised to a fixed breathing pattern due to their spontaneous nature. Normal breathing trials followed the participants’ usual respiratory rhythm. For deep breaths, participants first took two normal breaths, then performed the manoeuvre, a sequence designed to minimise any confounding effects on cerebrospinal fluid (CSF) and blood flow such as baseline shifts in venous return or intracranial pressure, and motion artefacts associated with continuous deep breathing. To reliably elicit yawns, participants were shown video clips of people and animals yawning while in the MRI scanner, a process used successfully in other studies (Chan et al., 2017; Gallup et al., 2019; Diana et al., 2023). For stifled yawns, participants were instructed to resist the urge to yawn once a response was triggered. All manoeuvres were performed within a single MRI session, with each respiratory action acquired in separate consecutive scans and repeated at least three times. Not all participants were able to complete every respiratory manoeuvre successfully; therefore, each analysis included only participants who met the relevant performance and repeat-scan criteria, with the total number of participants for each analysis specified in the Statistics section.

### Spirometry measurements

Spirometry was used to measure airflow volume during each manoeuvre outside the MRI, allowing correlation with MRI-derived CSF and cervical blood flow data. Airflow was recorded using a pneumotachometer (Series 3813; Hans Rudolph, Kansas City, KS) connected to a low-resistance face mask (Hans Rudolph) rather than a mouthpiece, enabling natural jaw and orofacial motion during yawning and deep-breathing tasks. The face mask configuration avoided the need for a mouthpiece, which can constrain jaw opening and alter orofacial muscle recruitment, thereby ensuring that the measured airflow more closely reflected the natural respiratory mechanics of yawning. Participants performed the same manoeuvres as during the MRI scans (normal breathing, deep breaths that mimic a yawn, yawning, and attempts to suppress yawns).

### MRI scans

MRI data were collected with a 3T Philips Ingenia CX scanner using a 16-channel neurovascular head and neck coil. Subjects underwent MRI scans to quantify orofacial movement, CSF, venous and arterial blood flow in response to the respiratory manoeuvres. Respiration was monitored concurrently by placing the vendor-supplied respiratory bellow the sternum, which tracked thorax displacement throughout the scans.

Real-time (RT) sagittal cine MRI and phase contrast MRI techniques were used to obtain time-courses of tissue movement and fluid velocities, respectively. Sagittal images were collected in the mid-sagittal plane using T1 fast field echo (T1FFE). Scanning parameters included: matrix = 112×112, FOV = 220mm, TR/TE = 4.1/2.3ms and a slice thickness of 10 mm. Axial phase contrast images were obtained at the level of the C3 vertebra, perpendicular to the spinal canal (Figure 1). At C3 both the CSF and blood flows through the plane and are within the field of view (FOV). The real-time PC-MRI protocol used turbo-field echo-planar imaging (TFEPI) and was not cardiac gated. Scanning parameters included: flip angle = 20°, matrix = 128×128mm, FOV = 192mm, repetition time (TR)/echo time (TE) = 13/7ms, and a slice thickness of 10mm. The encoding velocities were set to 10-20cm/s for CSF flow measurements and 60-90cm/s for arterial and internal jugular flow. The study collected data across 200 phases, with intervals of 70ms and 140ms. Whenever aliasing artifacts appeared, scans were conducted again using a higher encoded velocity to minimize these errors.

**Figure 1.**
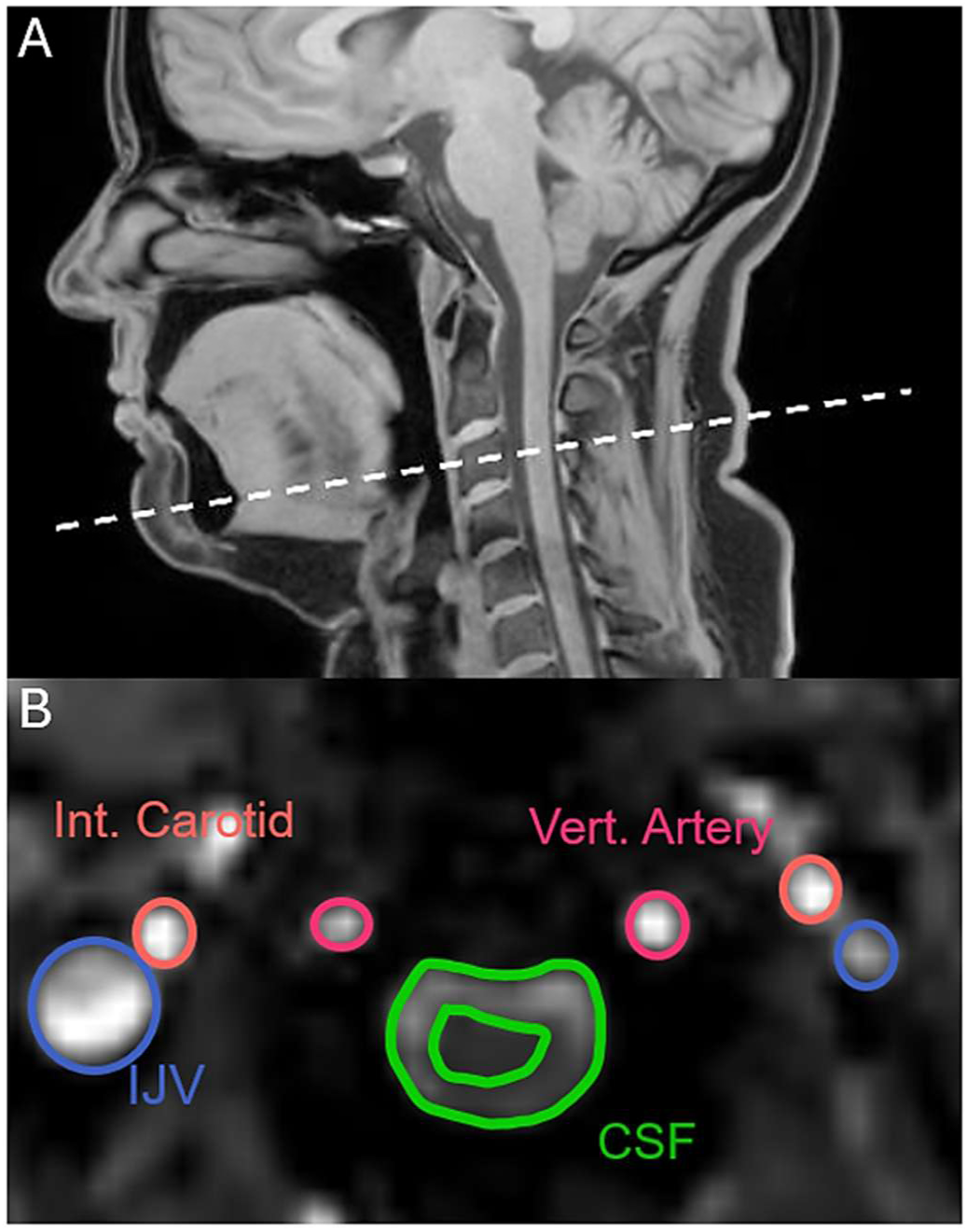
Imaging planes and analysis regions of interest. **(A)** Mid sagittal scan with the C3 imaging plane indicated with dashed white line. **(B)** Cropped view of the imaging plane at mid-C3 showing regions of interest: CSF (between green lines), internal jugular vein (blue), internal carotid (orange), and vertebral arteries (pink).

### Peripheral nerve stimulation scoring

Peripheral nerve stimulation (PNS) induced by rapidly switching magnetic field gradients during MRI can activate skeletal muscles, particularly in the thoracic and abdominal regions (Formica et al., 2004). Such stimulation can elicit brief or sustained contractions of intercostal, abdominal, or shoulder muscles, potentially influencing respiratory effort and the accuracy of voluntary breathing manoeuvres. To assess the presence and severity of PNS during scanning, participants self-reported any sensations after each scanning session using a five-point scale: 1 = no sensation, 2 = slight sensation, 3 = consistent stimulation throughout the scan, 4 = consistent and uncomfortable stimulation, and 5 = stimulation requiring early termination of the experiment.

### Data analysis

#### ROI segmentation and flow velocity extraction

Regions of interest (ROIs) comprised the cerebrospinal fluid (CSF) space, internal jugular vein (IJV), internal carotid artery, and vertebral artery at the C3 vertebra. For each manoeuvre, up to three MRI datasets were manually annotated to generate ground-truth masks. These labels were used to train nnU-Net, an out-of-the-box, self-configuring segmentation framework that automates preprocessing, network design, and training hyperparameters (Isensee et al., Nat Methods, 2021). We used the standard 3D configuration with five-fold cross-validation and commonplace augmentations (random flips, elastic deformations, intensity scaling). The model architecture and training approach were based on a previously published framework (Lloyd et al., 2025), in which nnU-Net models were trained for multiple cranio-cervical regions, including mid-C3 PC-MRI data. For each model, approximately 20 % of the training data were withheld for validation; the mid-C3 model was trained on 26 scans and validated against 10 withheld scans. The same trained mid-C3 model was used here without modification. Further details of that validation and model availability are provided in the Methods of Lloyd et al. (2025) and the trained models are available at: http://doi.org/10.17605/OSF.IO/HTE5Y, which can be used with 3D slicer by other researchers.

The trained model produced per-frame segmentation masks that were converted to region-of-interest (ROI) contours, representing the boundaries of the segmented structures in each frame. These contours were imported into Segment software (Heiberg et al., 2010), which uses them to define spatial masks for quantitative analysis. Within each ROI, Segment extracts pixel-wise velocity information from the corresponding phase-contrast MRI data, enabling computation of mean flow rates and temporal flow waveforms across the ROIs. Phase-contrast data occasionally exhibited residual phase wrapping, which was corrected manually by adding or subtracting 2π from affected pixel values (Bioucas-Dias & Valadao, 2007; Lloyd et al., 2020). CSF and blood velocities were extracted from the ROIs in Segment, and instantaneous flow rates were computed by multiplying velocity by ROI area.

CSF, blood, and respiratory air flow data were partitioned into epochs corresponding to each respiratory manoeuvre (normal breathing, deep breaths, yawns, and stifled yawns) using the respiratory band signal to identify event timing. Peak cranial and caudal CSF velocities and the net flow direction were compared across manoeuvres; equivalent analyses were performed for venous and arterial blood flow. Extracted flow and velocity metrics were subsequently used for within-subject and between-group statistical comparisons, as detailed below.

### Tongue motion tracking

Tongue motion was quantified using image registration to derive deformation fields between consecutive sagittal MRI frames. MATLAB’s *imregdemons* function (R2021a) was used to generate a pixel-wise deformation field that corrects for image distortions, rotations, and translations by aligning each moving frame to a stationary reference. A landmark on the tongue blade, just posterior to the tip, was selected because this region exhibited pronounced displacement during yawning while maintaining more consistent tracking than the tip itself. The tongue blade also provided clearer tissue contrast and a stable submucosal fat pocket that improved the reliability of marker placement across frames. Tongue positions were initialised manually at three points and automatically updated across frames using the deformation vectors.

Tongue motion similarity across scans was quantified using the MATLAB function corr, which computes the Pearson correlation coefficient (r) to assess linear association between variables (r = 1, perfect; r = 0, none; r = –1, inverse). Cross-correlation was selected because it quantifies the temporal alignment and waveform similarity of motion trajectories independent of absolute displacement magnitude. Correlations were calculated separately for x- (anteroposterior) and y- (superoinferior) displacements. To obtain a single measure of two-dimensional trajectory similarity, mean-centred x and y displacements were combined into displacement vectors, and a vector correlation coefficient was derived from their normalized dot product.

### Statistical analysis

All analyses were performed in GraphPad Prism 10.2.0 (GraphPad Software, Boston, MA, USA). Within-subject differences across respiratory manoeuvres (normal breathing, deep breaths, and yawns) were evaluated using one-way repeated-measures ANOVA. Between-group comparisons (e.g. sex or PNS scores) were assessed with two-tailed Mann–Whitney U tests. Data are presented as mean ± SD, and statistical significance was set at α = 0.05 (two-tailed). Significance levels are denoted as *** p < 0.001**, ** p < 0.01**, * p < 0.05**, and ns = not significant.

Participant inclusion varied across analyses according to task-performance criteria. All participants (n = 22) provided peripheral nerve stimulation (PNS) scores, whereas RT-PCMRI (n = 11) and spirometry flow comparisons (n = 13) included only those who completed all respiratory manoeuvres with at least two repeat scans per condition. Yawning-similarity analyses included participants with at least two successfully recorded yawns during sagittal real-time imaging (n = 18).

## Results

### Flow dynamics during normal breathing, gaping deep breaths, and yawning

MRI scan data from a representative participant demonstrates the dynamics of CSF and venous blood flow during normal breathing, deep breathing, and yawning episodes (Figure 2). During normal breathing, CSF and venous blood flow exhibited low-amplitude, low-frequency oscillations coupled to the respiratory and cardiac cycles, with minimal net directional change across the breathing cycle. During deep breaths, flow amplitudes increased substantially. Upon deep inspiration, CSF flow was directed cranially (towards the brain), while expiration produced a caudal (toward the spinal canal) shift. Venous return via the IJV also increased during inspiration, corresponding to increased outflow from the cranium to the thorax, whereas expiration reduced venous return as CSF was displaced caudally (Figure 2 and Table 1). Arterial inflow via the carotid and vertebral arteries increased transiently during inspiration. During yawning, changes in fluid flow were largest during the initial sharp inspiration, “gaping”, and expiratory phases. To best explain the results, the mechanics are separated into the four periods of yawning shown in Figures 2 and 6. For this representative dataset, the observations in each phase were:

i. Onset. A brief inspiration with no change in neurofluid flows compared to normal breathing.
ii. Maximal mouth opening with sharp inspiration.

a. Internal jugular vein (IJV) flow increased in magnitude, rising from a mean −7.5 mL/s to -13.6 mL/s; pulsation amplitude decreased, consistent with sustained venous outflow.
b. CSF simultaneously flowed caudally, concurrent with venous drainage.
c. The net CSF flow magnitude decreased slightly.
iii. Sharp expiration with mouth closing.

a. Cranial CSF flow increased during expiration compared with normal breathing.
b. During the brief gaping phase and early expiration, internal carotid arterial flow increased by ∼34%, whereas vertebral arterial flow remained near baseline.
iv. Recovery. A brief post-expiratory pause marked the end of the manoeuvre, followed by normalization of neurofluid flow. Yawning produced transient cardiovascular changes, including increased heart rate (as also reported by Heusner, 1946 and Askenasy, 1996), and often concluded with a swallow that can trigger a short-lived sympathetic response and heart rate acceleration (Burke et al., 2020).

**Figure 2.**
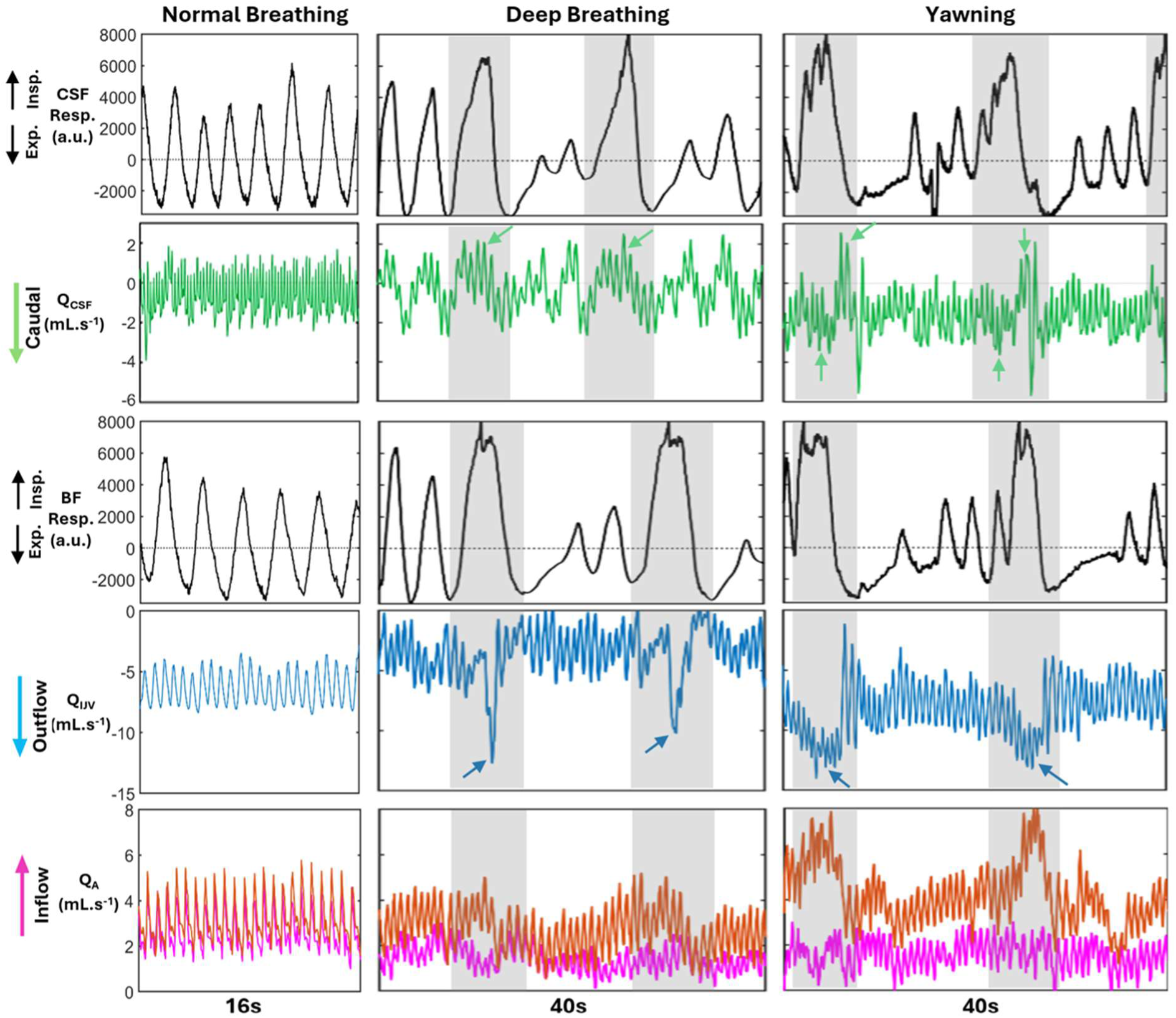
Respiratory-linked cerebrospinal fluid (CSF) and blood flow dynamics in a representative participant during normal breathing, deep breathing, and yawning. Panels show CSF (green) and vascular flows (Q in ml.s^-1^) from the internal jugular vein (blue), vertebral artery (orange), and internal carotid artery (magenta); positive values denote cranial flow and negative values caudal. Respiration (black trace, arbitrary units) is shown above each flow plot. Arrows mark CSF and IJV peaks. During normal breathing, CSF and venous flows show low-amplitude oscillations coupled to respiration and cardiac pulsation. Deep breathing produces larger bidirectional shifts relative to normal breathing, with cranial CSF inflow and increased jugular outflow on inspiration, reversing on expiration, consistent with pressure-driven coupling between thoracic and cranial compartments. In this participant, yawning elicited the largest changes in both CSF and blood flow, with CSF and venous outflow frequently aligned caudally during inspiration. Arterial inflow showed a brief inspiratory rise, particularly in the carotid trace. Velocity encoding was optimised separately for CSF and blood. CSF and blood flow scans were acquired sequentially, so respiratory traces are not time-locked across panels.

**Table 1.** Mean peak cerebrospinal fluid (CSF) and internal jugular vein (IJV) flow during normal breathing, deep breaths, and yawns. Values are mean ± SD across all participants. For CSF positive (cranial) and negative (caudal) values denote flow direction relative to the cranio-spinal axis. For the IJV, caudal values represent venous outflow from the cranium and cranial is the minimum outflow.

| | CSF (Mean ± SD) | | | IJV (Mean ± SD) | | |
| --- | --- | --- | --- | --- | --- | --- |
|  | Normal breathing | Deep breaths | Yawns | Normal breathing | Deep breaths | Yawns |
| **Cranial flow (mL/s)** | 1.86 ± 0.64 | 2.65 ± 0.81 | 2.56 ± 0.8 | -0.89 ± 0.8 | -1.17 ± 0.91 | -1.1 ± 0.77 |
| **Caudal flow (mL/s)** | -3.43 ± 1.65 | -3.69 ± 1.46 | -3.49 ± 1.34 | -5.44 ± 2.62 | -8.34 ± 3.0 | -9.53 ± 3.6 |

Mean peak neurofluid flow magnitudes for cranial and caudal flows, calculated as the average of peak values per manoeuvre, are presented for each participant (Figure 3). Table 1 summarises these across all subjects. Analysis revealed a significant increase in maximal cranial CSF flow through the C3 vertebral region during deep breathing (ANOVA, p = 0.003) and yawning (ANOVA, p = 0.006), demonstrating a clear relationship between enhanced respiratory effort and increased CSF flow compared to normal breathing. One experimental goal was to replicate yawning-induced flow magnitudes using a gaping deep breath to determine if meaningful differences exist between the two activities. However, no significant difference in CSF flow magnitude was observed between yawning and deep breathing (ANOVA, p = 0.894). Additionally, caudal IJV flow increased during both deep breathing and yawning compared to normal breathing. Stifled yawns were omitted from quantitative analysis due to the wide variety of suppression tactics used (complete or partial mouth closure, breath-holds, nasal breathing) yielding CSF and IJV flow profiles too variable for dependable direction or magnitude quantification.

**Figure 3:**
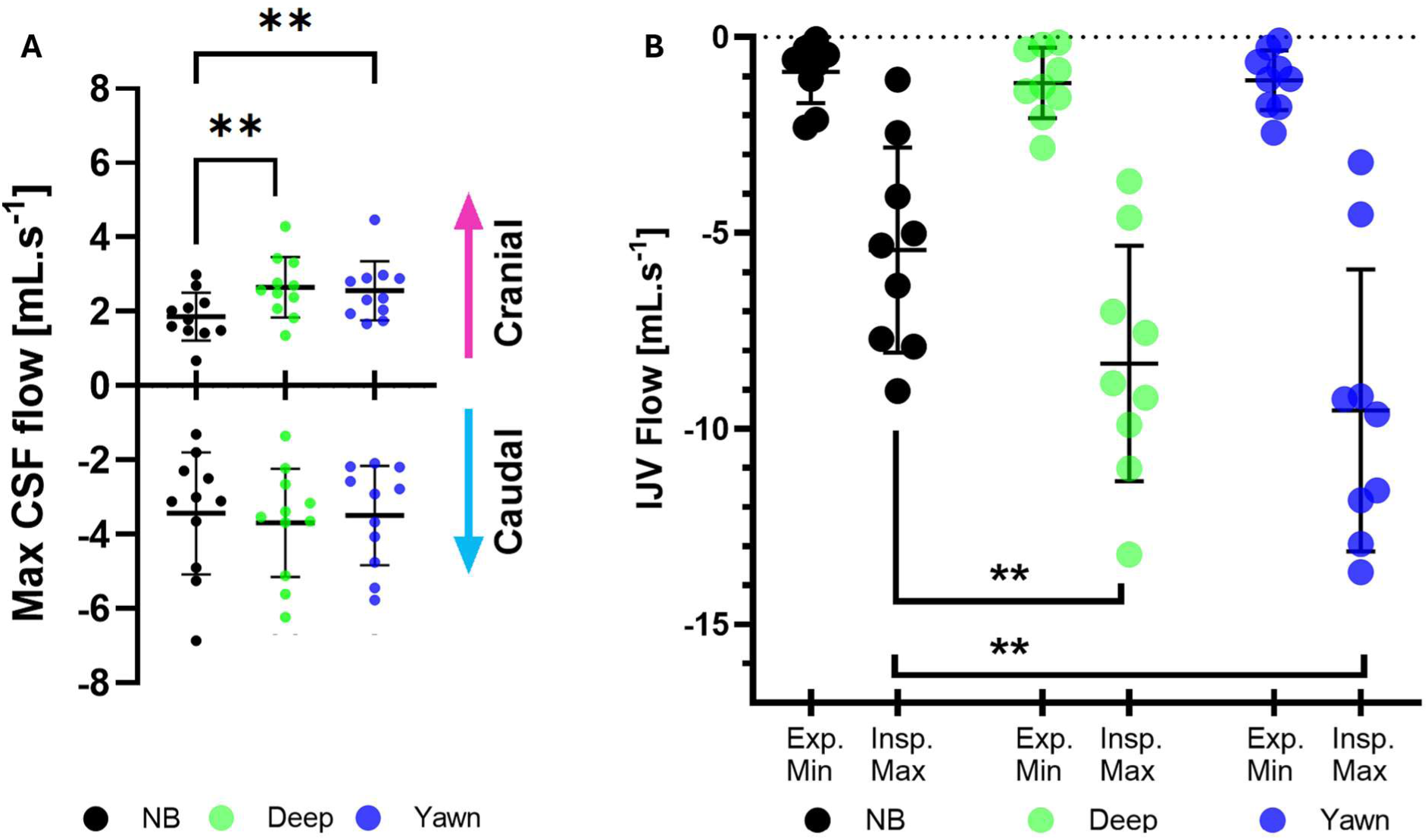
G**r**oup **data showing maximum and minimum cerebrospinal fluid (CSF) and internal jugular vein (IJV) blood flow at the C3 vertebra during normal breathing (NB), deep breaths, and yawning**. **(A)** Cranial CSF flow significantly increased during deep breathing (**p = 0.0032) and yawning (**p = 0.0058) compared with normal breathing. The study aimed to determine whether deep gaping breaths could replicate yawning-induced flow magnitudes; results showed no significant difference between the two manoeuvres (p = 0.8943). Caudal CSF flow remained consistent across all respiratory manoeuvres. **(B)** Minimum IJV blood flow was not different between normal breathing, deep breaths, and yawning. However, maximum IJV significantly increased during both deep breathing (**p = 0.007) and yawning (p = 0.01), indicating that enhanced respiratory effort boosts caudal IJV outflow. Statistical comparisons were performed using a one-way repeated measures ANOVA (** indicates p < 0.01; ns = not significant).

### Deep breathing and yawning neurofluid flow direction

At C3, inspiratory events were classified as *co-directional* when IJV and CSF flows were both caudal (co-outflow from the cranium), and *counter-directional* when CSF flow was cranial during maximal IJV outflow. Expiratory events invert these polarities by definition; our classification is referenced to inspiration. Counter-directional CSF and IJV flows were observed predominantly during normal respiration and deep breathing. During gaping deep breaths, all participants showed a consistent pattern of increased IJV outflow during inspiration, irrespective of sex. Similarly, yawning was consistently associated with increased IJV outflow during inspiration across all participants, reinforcing the observation of consistent venous flow patterns in response to both deep breaths and yawning. Unlike normal and deep respiration, which typically produced counter-directional CSF–IJV flow, yawning exhibited greater variability. In many instances of yawning, CSF and IJV flows were co-directional (Figure 4B, C), suggesting that yawns may modulate CSF–venous coupling differently from other respiratory manoeuvres. Also, a sex-specific pattern emerged: males predominantly exhibited counter-directional flow during yawns (76%), whereas females more often showed co-directional flow (74%) (Figure 4A, C), indicating possible sex-related variability in CSF dynamics. However, peripheral nerve stimulation (PNS) during PC-MRI was scored as strong by male participants. Several males described intense abdominal sensations, and in one case involuntary breathing was triggered, whereas most females reported no such stimulation. On the post-scan PNS questionnaire, female participants had a mean PNS score of 1.46 ± 0.69 (almost no sensation), significantly lower than the male average of 3.17 ± 1.03 (consistent stimulation throughout the scans) (Mann–Whitney U test, p = 0.0002; n = 11 females, n = 12 males). As PNS was rated once at the end of the scan session, scores reflect overall rather than manoeuvre-specific experiences (see Discussion/Limitations).

**Figure 4.**
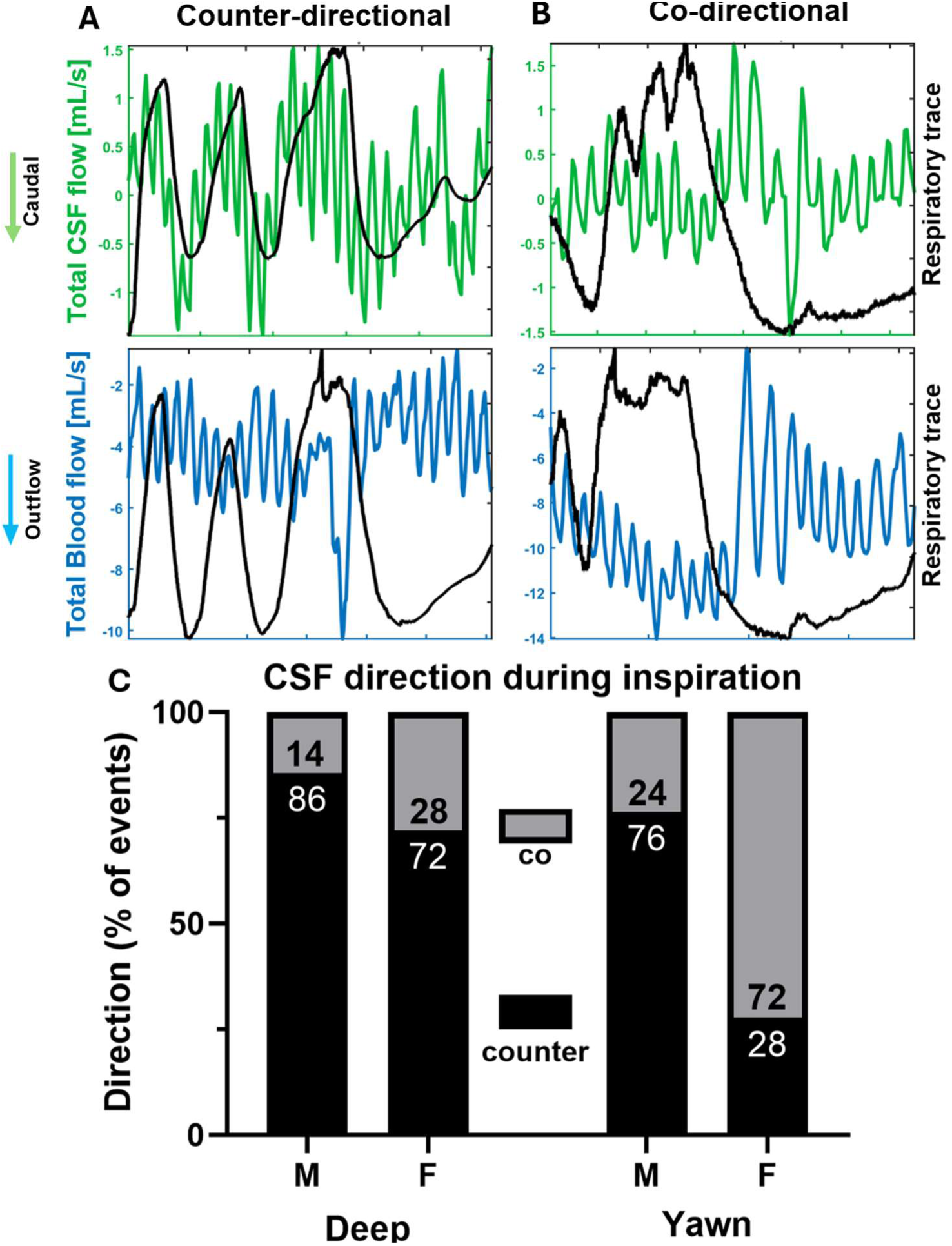
CSF-IJV directionality and synchrony during deep breathing and yawning. **(A)** Example of a counter-directional pattern, where cranial CSF flow (green) coincides with caudal venous outflow (blue) during inspiration (black respiratory trace). **(B)** Example of a co-directional pattern, where both CSF and venous blood flow caudally during inspiration. Positive values denote cranial flow; negative values denote caudal (outflow) direction. **(C)** Group data showing the proportion of counter- versus co-directional CSF–venous flow events during inspiration, separated by sex (M, male; F, female) and manoeuvre (deep breathing, yawning). During deep breathing, CSF and venous flows were predominantly counter-directional (males = 18/21 events ≈ 86%; females = 13/18 events ≈ 72%). Yawning also showed mainly counter-directional flow in males (13/17 events ≈ 76%) but a much greater proportion of co-directional flow in females (13/18 events ≈ 72%). Possible effects of peripheral nerve stimulation (PNS) on male yawning data are discussed in the Limitations.

### Spirometry measurements for deep breathing, yawning, and stifled yawns

Spirometry data from a representative participant illustrate airflow differences between deep breaths, yawns, and stifled yawns (Figure 5A). Mean peak airflow for normal breathing inspiration was 0.54 ± 0.13 L/s, deep breath inspiration was 0.98 ± 0.39 L/s, and yawning inspiration was 1.03 ± 0.52. L/s. For expiration, the peak airflow was −0.49 ± 0.13 L/s for normal breathing, −0.99 ± 0.45 L/s for deep breaths, and −1.41 ± 0.69 L/s for yawns. (Figure 5B). Both deep breaths and yawns exhibited greater inspiratory and expiratory airflow compared with normal breathing. However, the difference in airflow between deep breaths and yawns were not significant (Inspiration: p-value of 0.946; Expiration: p-value of 0.077). Swallowing occurred within one breath most commonly after yawning (81 ± 14%), slightly less common after a stifled yawn (68 ± 8.4%), and less common after a deep breath (16 ± 23%).

**Figure 5.**
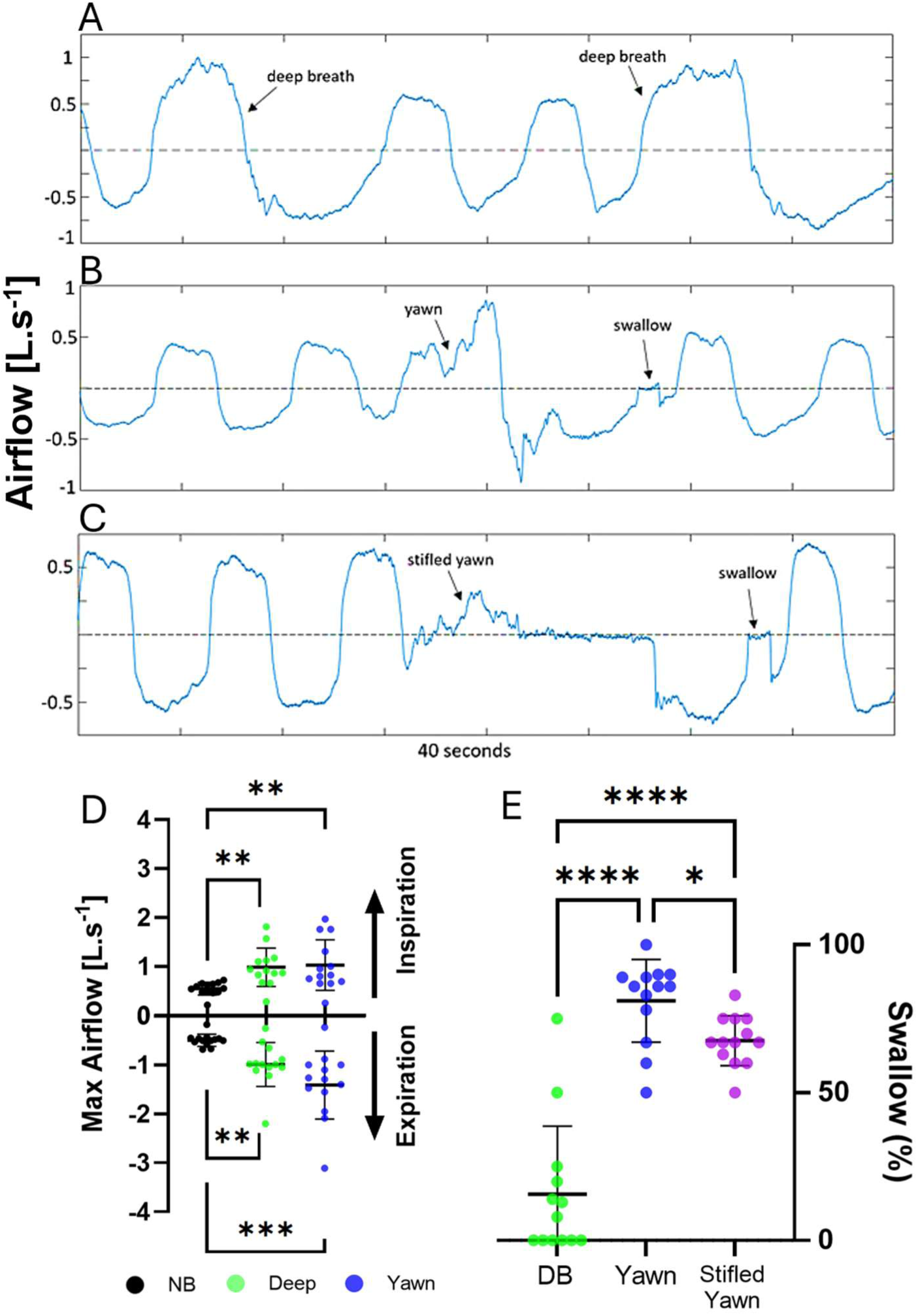
Spirometry during deep breaths, yawns, and stifled yawns (representative participant) with group airflow statistics. **(A)** Deep-breath airflow trace; arrows indicate two gaping deep breaths. **(B)** Yawn airflow trace; arrows mark a yawn and a swallow. **(C)** Stifled-yawn airflow trace; arrow marks a swallow. Airflow for stifled yawns is not reported because rapid mouth closure to suppress the yawn produced incomplete measurements. **(D)** Proportion of manoeuvres (deep breaths, yawns, stifled yawns) followed by a swallow within one breath (∼5 s). Peak airflow (mean ± SD). Inspiration – normal breathing: 0.54 ± 0.13; deep breath: 0.98 ± 0.39; yawn: 1.03 ± 0.52. Expiration – normal breathing: −0.49 ± 0.13; deep breath: −0.99 ± 0.45; yawn: −1.41 ± 0.69. Deep breaths and yawns exceeded normal breathing (Inspiration: p = 0.0016 and 0.0066; Expiration: p = 0.0010 and 0.0007). Deep breaths vs yawns: not significant (Inspiration: p = 0.9463; Expiration: p = 0.0773). **(E)** 81 ± 14% of yawns and 68 ± 8.4% of stifled yawns, and 16 ± 23% of deep breaths were followed by a swallow within one normal breath (approximately 5 s).

**Figure 6.**
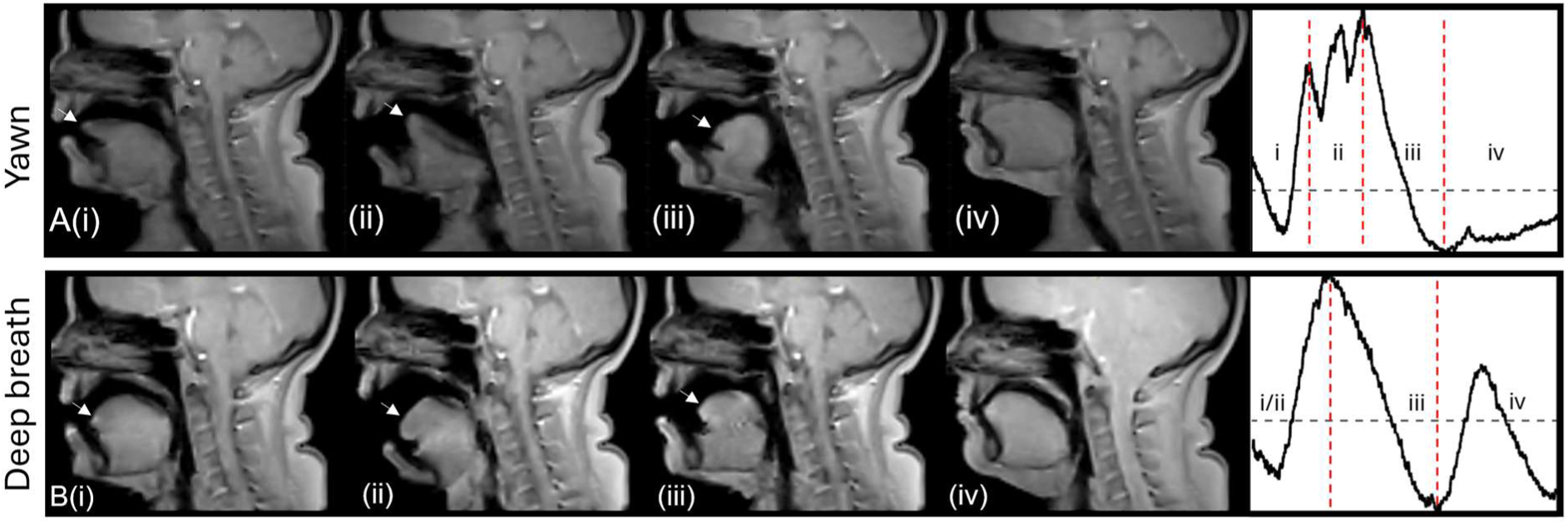
Sequential sagittal MRI frames showing tongue and oropharyngeal motion during yawning and deep breathing with a gape. **(A)** Yawning timelapse (Video 1): sagittal frames (i–iv) illustrate key phases of (i) onset, tongue retracts, inspiration begins (ii) maximal mouth opening with sharp inspiration, (iii) sharp expiration with mouth closing; tongue shape may change, and (iv) recovery. The corresponding respiratory trace (right) indicates each phase (red dashed lines). **(B)** Timelapse of a deep breath with a gape (Video 2): sagittal frames (i–iv) show a similar inspiratory–expiratory sequence but with reduced orofacial displacement compared to yawning. (i) inspiration with mouth opening; (ii) mouth opens further as inspiration progresses; tongue position/shape remains relatively unchanged; (iii) expiration with mouth closing. The corresponding respiratory trace (right) indicates each phase (red dashed lines). White arrows denote the tongue tip position in each frame. Images were acquired in real time at the mid-sagittal plane. Positive respiratory deflection indicates inspiration.

### Sagittal real-time montage of yawn and deep breaths

The yawning cycle observed during this study consisted of four phases (Figure 6A): retraction of the tongue and sharp inspiration (i). immediately followed by a phase of sharp inspiration that coincides with the opening of the mouth. This gaping phase lasts approximately one second and is the apex of the yawning process (ii). The gaping phase transitions into a sharp expiration, during which the mouth begins to close. It is noteworthy that during this phase, alterations in the shape of the tongue were observed in most participants (iii). The cycle concludes with the participant returning to a resting state, characterised by a return to normal breathing patterns (iv). For comparison, a montage of a deep breath is shown in Figure 6B. Many features of the gaping deep breath are similar to the yawn; however, yawning is distinguished by more pronounced tongue motion and deformation.

### Kinematics of the tongue during yawning

During the investigation of yawning effects on CSF and blood flow, we found that yawning kinematics, while following a common sequence (Figure 7), were distinct between individuals. Once triggered, yawns proved difficult to suppress completely. While participants could modify the yawn by keeping their lips closed and reducing jaw movement, they could not prevent the yawn entirely. In the example in Figure 7, an unstifled yawn (A) is compared with a stifled yawn (B); despite the stifling, both exhibit the characteristic tongue shape after initiation (frames 2–4) and the rapid horizontal-to-vertical “flip” at the peak. Across the cohort, tongue trajectories were tracked for all participants; despite inter-participant variation, each participant’s repeated yawns were similar (mean intra-participant cross-correlation 0.86 ± 0.062), and in participants who did not modify their yawns, successive trajectories were near-identical (Ci, ii) often differing by only one to two pixels.

**Figure 7.**
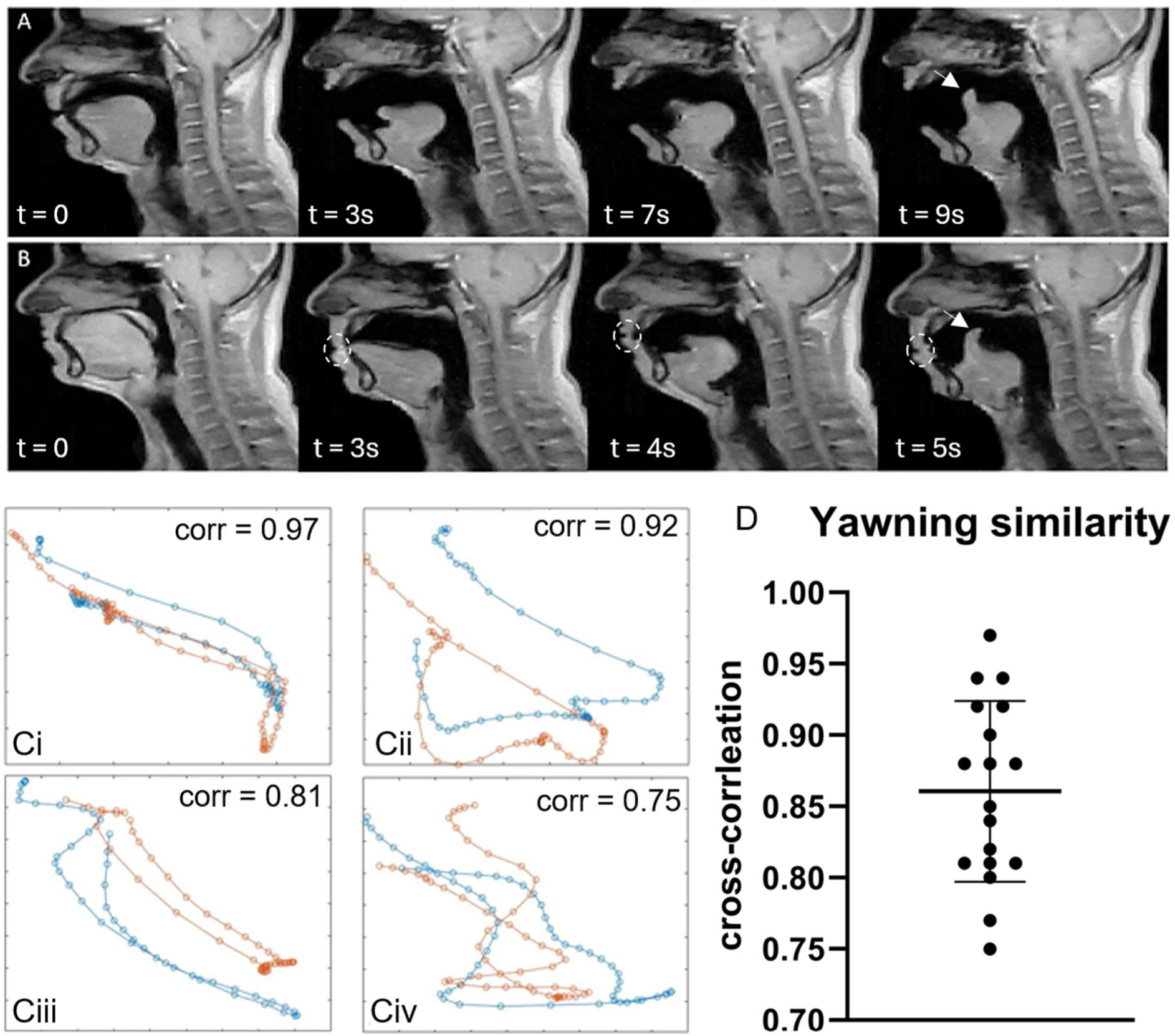
Sagittal real-time scans of a yawn and a stifled yawn in the same individual and tongue motion similarity across the cohort during yawning. (A) Montage of a complete yawn (Video 3): the tongue retracts after inspiration, then rapidly alternates between horizontal and vertical orientations at the yawn’s apex (white arrows), before expiration and return to rest. (B) Montage of a stifled yawn (Video 4): despite the closed mouth, the tongue movements are similar to the full yawn. Dashed white circle shows that the mouth was voluntarily kept closed. (C) Cross-correlation analysis. Tongue trajectories and similarity coefficients for four participants. X and Y axes represent sagittal scan pixel positions; orange and blue traces denote different yawns. Each participant shows a distinct yawning pattern which is similar within individuals but distinct across participants. The cross-correlation coefficient (corr), calculated as described in the Methods, averaged (D) 0.86 ± 0.062 across participants. An example of the tracking is given in Video 5).

## Discussion

Although respiration is a dominant driver of CSF movement, the effects of yawning on CSF and blood flow have not been previously characterised. In this study, we show that yawns and deep breaths produce comparably large CSF and venous flow magnitudes to normal breathing, but they differ in flow directionality. Deep inspirations generally increased cranial CSF flow with counter-directional IJV outflow, whereas yawns more often produced co-directional caudal CSF and IJV flows during inspiration. In addition, yawns within individuals also exhibited highly stereotyped orofacial tongue kinematics across repeated events, compatible with, but not proof of, control by a central pattern generator. This has not been previously identified.

### Comparison of neurofluid flow between deep breathing and yawning

The magnitude of CSF flow during deep breaths and yawns was found to be similar, and both were greater than normal breathing. These findings align with the spirometry results, which also showed comparable airflow rates for yawning and deep breathing. During deep inspiration, there was an increase in cranial CSF flow, while expiration prompted a caudal movement of CSF. The IJV flow magnitude during deep breaths were greater than in normal breathing and flowed in the counter-directionally to CSF. These patterns align with prior reports that forced breathing increases CSF flow compared with eupnoea and introduces physiological variability across sites (Yamada et al., 2013; Dreha-Kulaczewski et al., 2015, 2018; Gutiérrez-Montes et al., 2022; Kollmeier et al., 2022). However, during yawning, CSF flow direction differed from both normal and deep inspiration.

Venous–CSF coupling during deep breathing likely reflects changes in intrathoracic and intra-abdominal pressures (Lloyd et al., 2020). In our data, caudal IJV flow during inspiration was consistent across participants for both gaping deep breaths and yawns. In contrast, CSF directionality during yawning differed from deep breaths: despite similar peak magnitudes, yawns frequently produced co-directional caudal CSF and IJV flows (Figures 2, 4). Pressure differences between the cranium and thorax are the driving force for CSF and venous blood flow during respiration yet direct measurements of intrathoracic/cervical pressures during human yawns are not available. However, a plausible mechanism to explain the differences between deep breathing and yawning is that ribcage expansion and diaphragmatic descent generate negative intrathoracic pressure (enhancing venous return) (Agostini & Rahn, 1960; Kono & Mead, 1967), while pharyngo-laryngeal dilation, mediated by activation of upper-airway dilator muscles, can lower upper-airway resistance (Pierce et al., 2007). Together with orofacial–pharyngeal muscle recruitment, this may transiently bias CSF caudally during yawning (Lloyd et al., 2025). This remains hypothetical and warrants simultaneous pressure–flow recordings in future studies.

### Sex Differences in CSF flow direction during yawns and peripheral nerve stimulation

Analysis of CSF flow during yawning revealed possible sex-specific differences (Figure 4C). In males, CSF and IJV flows were typically counter-directional, whereas females more often exhibited co-directional flows. All males reported strong PNS sensation while 7/11 females reported none. This PNS induced abdominal activation likely increased intrabdominal pressures and restricted diaphragmatic excursion, and disrupted normal breathing coordination, potentially influencing CSF flow directionality in males (Agostini and Rahn., 1960; Lloyd et al. 2020, 2025). Consequently, female data in this study may better reflect normal yawning physiology outside the MRI environment (see Limitations).

### Evidence that yawning is controlled by a central pattern generator (CPG)

Despite inter-individual variation, consistent stereotyped behaviours were observed across multiple yawns for a given individual (Figure 7). Intra-individual similarity in yawning patterns was high, with cross-correlation coefficients ranging from 0.75 to 0.97 for tracked tongue movements (group mean was 0.86 ±0.062). Even when yawns were stifled, the tongue exhibited movements quite like those seen in full yawns, indicating that the yawning motor pattern, once initiated, is automatic and difficult to suppress or alter (Figure 7). This balance between intra-subject consistency and inter-subject variability indicates that yawning displays individual-specific motor patterns, consistent with, but not definitive evidence of, a yawning CPG. CPGs operate autonomously, generating patterns of neural activity that drive rhythmic behaviours (Dzeladini et al., 2014; Katz., 2016; Krestel et al., 2018). In the case of yawning, the CPG would autonomously initiate and execute the yawning cycle, explaining the consistent intra-individual flow and tongue movement pattern (Walusinski., 2010). Despite their autonomous nature, CPG outputs can be modulated by external stimuli or internal states (Straub et al., 2024). This flexibility might account for the variations in inter-participant yawning patterns while still maintaining a recognisable, individual-specific pattern; and implies that the patterns of yawning are not learned but are an innate aspect of neurological programming.

This hypothesis is bolstered by observations (Figure 6) which show that in ∼81 ± 14% of induced yawns, and in 68 ± 8.4% of stifled yawns, a swallow followed within one breath (Figure 5), suggesting close functional coupling. Yawning and swallowing, though outwardly appearing as distinct physiological behaviours, may be closely interconnected through their underlying neurological mechanisms since spontaneous yawning is frequently associated with spontaneous swallows (Abe et al., 2015) and are hypothesised to be influenced by a network of brainstem regions that includes CPGs responsible for both behaviours (Ertekin et al., 2015). Swallowing is organised by a medullary CPG with a dorsal swallowing group in the nucleus tractus solitarius and a ventral swallowing group near the nucleus ambiguous (Jean, 2001; Ertekin & Aydogdu, 2003). This network interacts with the respiratory pattern generator to coordinate brief swallow apnoea and airway protection (Jean & Dallaporta, 2006; Bianchi & Gestreau, 2009). The frequent yawn–swallow pairing therefore likely reflects interacting pattern generators and shared orofacial–pharyngeal synergies, which may contribute to the reproducible and distinct CSF–IJV flow alignment observed in our data during yawning.

### Implications of directionality of CSF and blood flows during yawning: brain waste clearance and thermoregulation

#### Waste clearance

Yawning is a coordinated neuromuscular activity that impacts fluid dynamics in the cranial and cervical regions. On this basis, yawning has been proposed to facilitate brain waste clearance via the glymphatic system (or cerebral waste clearance generally) potentially by augmenting venous return and promoting transit of cervical lymphatic fluid into central vessels during neck flexion (Dolkart, 2017). However, to date there has been limited direct evidence for this. Deep breathing increases intracranial arterial and venous volume displacement (Burman & Alperin, 2024), supporting a respiratory mechanism; our data extend this by showing that yawning, while comparable in magnitude to deep breathing, differs in CSF–IJV coupling, with frequent co-directional caudal flows during yawning inspiration. We interpret these as transient, jointly directed outflow that could favour caudal advection toward the spinal canal and thereby enhance macroscopic CSF mixing or clearance, particularly around sleep–wake transitions (Guggisberg et al., 2007; Zilli et al., 2007). However, this remains an indirect inference: whether such changes in velocity and directionality translate into greater parenchymal waste removal has not been demonstrated in humans and needs to be evaluated in future work.

#### Thermoregulation

The thermoregulatory hypothesis for yawning suggests that yawning helps dissipate excess brain heat by increasing airflow and heat exchange (Gallup and Eldakar., 2013) We found that both yawning, and deep breathing increased CSF and blood circulation compared with normal breathing. Furthermore, the co-directional flow of CSF and venous return could increase heat transfer from the brain to the lungs. This coordinated respiratory and vascular response during yawning appears optimized for maximal fluid exchange, producing the largest combined displacement of venous blood and CSF of any spontaneous respiratory manoeuvre. The human brain operates at a higher temperature (0.3°C to 0.93°C ± 0.5°C) than the body’s core (Mcilvoy, L., 2004; Oh et al., 2020). The alignment of CSF and venous blood flow could facilitate increased (compared to normal and deep breathing) heat transfer during inspiration where hotter CSF and venous blood leave the brain, while arterial flow from the thorax would cool the brain. This process could not only enhance thermal regulation but also adheres to the Monroe-Kellie doctrine, which posits that the volume inside the cranium is fixed; thus, any increase in one component requires a decrease in another, maximising the capacity for heat exchange through blood and CSF.

Evidence from behavioural studies supports a thermoregulatory role for yawning through a hypothesized thermal window effect: yawning frequency increases when ambient temperatures are below body temperature but declines when conditions approach or exceed it (Gallup & Gallup, 2007, 2012; Corey et al., 2012; Gallup & Eldakar, 2013; Marraffa et al., 2017; Massen et al., 2021). This pattern reflects that cooling efficiency is greatest when inspired air and circulating blood can absorb excess heat, and least when ambient temperatures are too high for heat exchange. Experimental studies found that applying neck and facial cooling over the carotid regions, which facilitates cranial heat loss, reduced spontaneous yawning frequency (Ramirez et al., 2019). Taken together, the alignment of CSF and venous outflow observed here, and previously by Klose et al. (1994), could be a physiological mechanism that augments the cooling potential of yawning beyond that of ordinary respiratory manoeuvres. These data situate yawning within broader homeostatic functions without implying a single purpose.

### Study Limitations

We identified possible sex differences in fluid flow during yawning, with female participants showing co-directional CSF and IJV flow, whereas males displayed counter-directional flow directions. This discrepancy may result from PNS, experienced predominantly and more strongly by males. Consequently, female data might better represent typical neurofluid physiology. PNS during MRI scans results from rapidly changing magnetic fields inducing electrical currents in peripheral nerves via electromagnetic induction. This can cause transient sensations such as twitching, tingling, or motor activation, with severity influenced by magnetic field strength, patient positioning, and individual anatomy (Faber et al., 2003). Anatomical modelling shows males generally have lower PNS thresholds due to larger thoracic dimensions, while thicker perineural fat deposits in females may raise thresholds by reducing the electric field at the nerve (Davids et al., 2019). Positioning within the scanner can further modulate PNS effects (Faber et al., 2003). Positions nearer to the scanner’s isocentre are associated with higher PNS thresholds and thus less PNS sensation. This could explain why PNS was common in our study, as the isocentre was positioned at the C3 vertebra in the neck, placing the participants’ thorax and abdomen farther from the scanner’s isocentre, potentially resulting in lower PNS thresholds.

CSF and blood flow were measured in separate, sequential scans rather than simultaneously because of velocity encoding constraints; using a higher velocity encoding setting can introduce noise in slower flows, potentially reducing measurement accuracy. However, given the intra-individual consistency, this is unlikely to undermine our main findings. Dual-encoding PC-MRI techniques could mitigate this issue in future studies. Quantitative airflow and intrathoracic/intra-abdominal pressures were not measured in the MRI due to equipment constraints and invasiveness, though existing deep-breathing data (Lloyd et al., 2020) may partially inform these results.

### Conclusions

This study reported CSF and blood flow changes during yawning in humans. The results demonstrate that there are noticeable changes in neurofluid flow when yawning compared with normal and deep breathing. Interestingly, the direction of CSF flow during yawning appears to differ from deep breaths, although the magnitude of the flow changes is similar. The simultaneous caudal movement of CSF and IJV blood during yawn inspiration could be the maximal transfer of solute-rich neurofluid to the spinal canal, via respiratory mechanics, possibly serving as a peripheral mechanism for waste clearance. Though this idea is speculative, it introduces an interesting avenue for understanding the physiological functions of yawning. However, co-directional caudal CSF and IJV flows during yawning may simply reflect the composite motor and pressure patterns unique to yawns; whether such coupling materially affects clearance is unknown. This alignment of CSF and venous blood flow during yawning may also optimise heat exchange, contributing to cooling of the brain. Additionally, our data suggest that individuals exhibit characteristic yawning patterns, consistent with the idea that yawning may be coordinated by a central pattern generator (CPG). Finally, yawning appears to be highly adaptive behaviour and further research into its physiological significance may prove fruitful for understanding CNS homeostasis.

## Acknowledgements

This research was produced in whole or part by UNSW Sydney researchers and is subject to the UNSW Intellectual property policy. It was also funded in whole or part by the National Health and Medical Research Council [APP2011940]. For the purposes of Open Access, the author has applied a Creative Commons Attribution CC-BY licence to any Author Accepted Manuscript (AAM) version arising from this submission. The authors acknowledge the facilities and scientific and technical assistance of NeuRA Imaging, a node of the National Imaging Facility, a National Collaborative Research Infrastructure Strategy (NCRIS) capability. Adam Martinac and this research was supported by the National Health and Medical Research Council (NHMRC) Ideas grant [APP2011940]. Robert Lloyd was also supported by a NHMRC Ideas grant [APP2011940]. Lynne Bilston was supported by a NHMRC Senior Research Fellowships [APP107793].

## Data availability

The nnU-Net training framework and model implementation used in this study are publicly available through the open-access repository described in our previous work (Lloyd et al., 2025 https://doi.org/10.17605/OSF.IO/HTE5Y). The trained mid-C3 segmentation model applied here can be run directly in 3D Slicer using the provided instructions and configuration files. The data that support the findings of this study are not publicly available due to restrictions of the approved ethics.

## Author Contributions

A.D.M., S.W., R.L., and L.E.B., conceived and designed the experiments. A.D.M. and S.W. performed the experiments. A.D.M., and R.L., performed the analysis. All authors contributed to the interpretation of the data and writing the paper. All authors approved the final version of the manuscript submitted for publication and agree to be accountable for all aspects of the work. All persons designated as authors qualify for authorship, and all who qualify for authorship are listed.

